# Granular Transcriptomic Signatures Derived from Independent Component Analysis of Bulk Nervous Tissue for Studying Labile Brain Physiologies

**DOI:** 10.1101/2020.01.01.892281

**Authors:** Zeid M Rusan, Michael P Cary, Roland J Bainton

## Abstract

Multicellular organisms employ concurrent gene regulatory programs to control development and physiology of cells and tissues. The *Drosophila melanogaster* model system has a remarkable history of revealing the genes and mechanisms underlying fundamental biology yet much remains unclear. In particular, brain xenobiotic protection and endobiotic regulatory systems that require transcriptional coordination across different cell types, operating in parallel with the primary nervous system and metabolic functions of each cell type, are still poorly understood. Here we use the unsupervised machine learning method independent component analysis (ICA) on majority fresh-frozen, bulk tissue microarrays to define biologically pertinent gene expression signatures which are sparse, i.e. each involving only a fraction of all fly genes. We optimize the gene expression signature definitions partly through repeated application of a stochastic ICA algorithm to a compendium of 3,346 microarrays from 221 experiments provided by the *Drosophila* research community. Our optimized ICA model of pan fly gene expression consists of 850 modules of co-regulated genes that map to tissue developmental stages, disease states, cell-autonomous pathways and presumably novel processes. Importantly, we show biologically relevant gene modules expressed at varying amplitudes in whole brain and isolated adult blood-brain barrier cell levels. Thus, whole tissue derived ICA transcriptional signatures that transcend single cell type boundaries provide a window into the transcriptional states of difficult to isolate cell ensembles maintaining delicate brain physiologies. We believe the fly ICA gene expression signatures set, by virtue of the success of ICA at inferring robust often low amplitude patterns across large datasets and the quality of the input samples, to be an important asset for analyzing compendium and newly generated microarray or RNA-seq expression datasets.

## Introduction

Uncovering the gene expression patterns of health and disease remains a monumental task in biology. In no other tissue is this challenge more difficult than in the central nervous system (CNS). The CNS is believed to be the most complex organ in terms of cell types and architecture, transcriptional programs and functional outputs (Hawrylycz et al., 2015; Kelley and Oldham, 2015). Genetics has been used to study the CNS and while several processes have been successfully interrogated, many others such as behavioral output systems remain poorly understood. In particular, endobiotic, xenobiotic and metabolic perturbations and fluctuations produce profound behavioral change, but their CNS genetic encodement is nebulous and regulatory compensations are ostensibly distributed across cell types (Hindle and Bainton, 2014). Thus classical pathway discovery through genetics has proved difficult for such physiologies and may never overcome the homeostatic and distributed-system obstacles to understanding gene function in complex tissues.

The high-throughput gene expression profiling revolution of the 21^st^ century has elevated the quality and quantity of molecular genetic signatures collected from biological samples. New methods such as massively parallel single cell RNA sequencing (RNA-seq) provide valuable information about cell type enriched gene transcript levels (Poirion et al., 2016). However, these cell type specific transcriptomes are not readily converted into the active cellular processes that have a transcriptional element. Parallel transcriptional programs, intrinsic and extrinsic noise, and RNA stability mechanisms affect cellular RNA content. Thus, accurate transcriptomes from purified cells in normal and disease states are mixtures of concurrent processes that require deconvolution (Brown and Celniker, 2015).

Discovering co-expressed genes that correlate with biological states provides a foundation for deciphering gene function. Analyses that span multiple gene expression experiments have revealed consistent patterns shared across experiments. These so-called “meta-analyses” take on many forms but all aim to accurately define groups of co-regulated genes by drawing on more information than what is provided by individual experiments. Hierarchical clustering analysis (HCA) is perhaps the most famous of these methods where experimental conditions and genes can be clustered based on gene-level correlations between experiments (Eisen et al., 1998). Another correlation-based method, weighted gene co-expression network analysis (WGCNA), looks for sets of genes of an expression compendium that have similar gene-to-gene correlation network topographies (Horvath and Dong, 2008). While HCA reveals modules of mutually exclusive sets of genes, WGCNA allows for genes to be reused in its modules. However, both HC and WGCNA find global signals trying to fit a model to all the samples, and so struggle to reveal more local signals enriched only in subsets of samples (Engreitz et al., 2010; Lee and Batzoglou, 2003). Thus, these types of analyses are an alternative to paired gene mutation-phenotype analyses for deducing gene importance in a process. But, for finding modules related to more subtle physiologies in large-scale gene expression data sets, other types of meta-analyses based on higher-order statistics are required.

Independent component analysis (ICA) has proven to be a powerful technique for uncovering hidden patterns from mixtures since its invention in the 1980’s (Comon, 1994; Hyvärinen and Oja, 2000). This data matrix decomposition method like others generates a set of basis vectors whose combination at different levels can reconstitute any sample of the original data compendium (Alter et al., 2000; Carmona-Saez et al., 2006; Holter et al., 2000; Lee and Seung, 1999). Unlike the popular matrix decomposition method, principal component analysis (PCA), ICA’s basis vectors are uncorrelated and maximally non-Gaussian, i.e. are maximally statistically independent. In effect, the ICA model of expression data yields relatively small groups of co-regulated gene modules compared to those produced by PCA (Reviewed in Kong et al., 2008). This property is more biological, matching the vast evidence that only small subsets of all genes are important in a given independently regulated process. ICA enforces statistical independence on its extracted patterns by maximizing higher-order statistics in the basis vectors and has outperformed PCA and other methods in enriching for biological transcriptomic signatures in several species ranging from yeast to human (Engreitz et al., 2010; Hori et al., 2001; Kong et al., 2008; Lee and Batzoglou, 2003; Liebermeister, 2002). Moreover, ICA is believed to be more sensitive for finding low amplitude patterns of gene expression, given enough input data amount and pertinent perturbation diversity (Engreitz et al., 2010; Lee and Batzoglou, 2003). Despite this success, ICA has not been used on a genetically tractable organism with a complex CNS such as *Drosophila melanogaster* to our knowledge. The combination of *Drosophila* genetic tools, evolutionarily conserved CNS properties and an accurate representation of transcriptional programs would improve the fly research model.

In this paper, we use ICA on a compendium of expression data produced by the *Drosophila* research community over the last 15 years to extract fundamental gene co-expression modules. We optimize a set of 850 modules mined from 3,346 published microarrays found in 221 experiments with a significant CNS presence. This diverse microarray data compendium provides the varying combinations of independent transcriptional signatures that ICA is able to disentangle. Modules map to many cell biologic and cellular processes, and seemingly identify new patterns of transcription. By treating the modules as a general model of fly expression variance we reinterpret CNS cell transcriptomes, which were not part of the input data compendium, in the context of modules. We find that multiple biologically relevant modules map strongly to the expression changes found between glial and neuronal cells, indicating that ICA modules provide a more granular view of fly cell transcriptomes. We discuss the utility of such modules for focusing gene function and regulation studies, and their refinement with ever-increasing data.

## Results

### Examples of data mining via ICA

Prior to using ICA for gene expression module discovery we demonstrated its ability to separate source signals from mixtures with an intuitive example (**Figure 1A**). First, we generated two thousand image-mixtures (*I*_1_–*I*_2000_) from combining grayscale images (vectors of pixel values, s_1_–s_4_) and Gaussian noise (s_noise_). Random integer coefficients (*k* = 1–5) were applied to s_1_–s_4_ pixel values prior to their summation to vary signal amplitudes in the mixture set, then unique random *k* = 10 × s_noise_ was added to each individual mixture. Second, we applied singular value decomposition (SVD) to the centered and scaled *I*_1_–*I*_2000_ mixture compendium achieving a PCA (Alter et al., 2000; Holter et al., 2000). We also applied ICA to *I*_1_–*I*_10_ (low data) and ICA to *I*_1_–*I*_2000_ (high data) using the FastICA algorithm (Hyvärinen and Oja, 1997; Marchini and Ripley, 2013). ICA on high data (bottom row, **Figure 1A**) clearly outperformed SVD on the same data (top row, first five singular vectors shown, **Figure 1A**) and ICA on low data (middle row, **Figure 1A**) in estimating the hidden source signals s_1_–s_4_. Estimated signals from ICA lack the true source signal pixel intensity information and instead describe relative pixel intensity relationships, i.e. pixels now have signed *weights* rather than grayscale intensity scalars. Thus, unsupervised ICA succeeds in estimating source signals from abundant samplings of mixtures but cannot recover the original scale and sign; pure white pixels may be estimated as pure black while relative intensity difference information is maintained in a different stochastic FastICA run.

**Figure 1.**
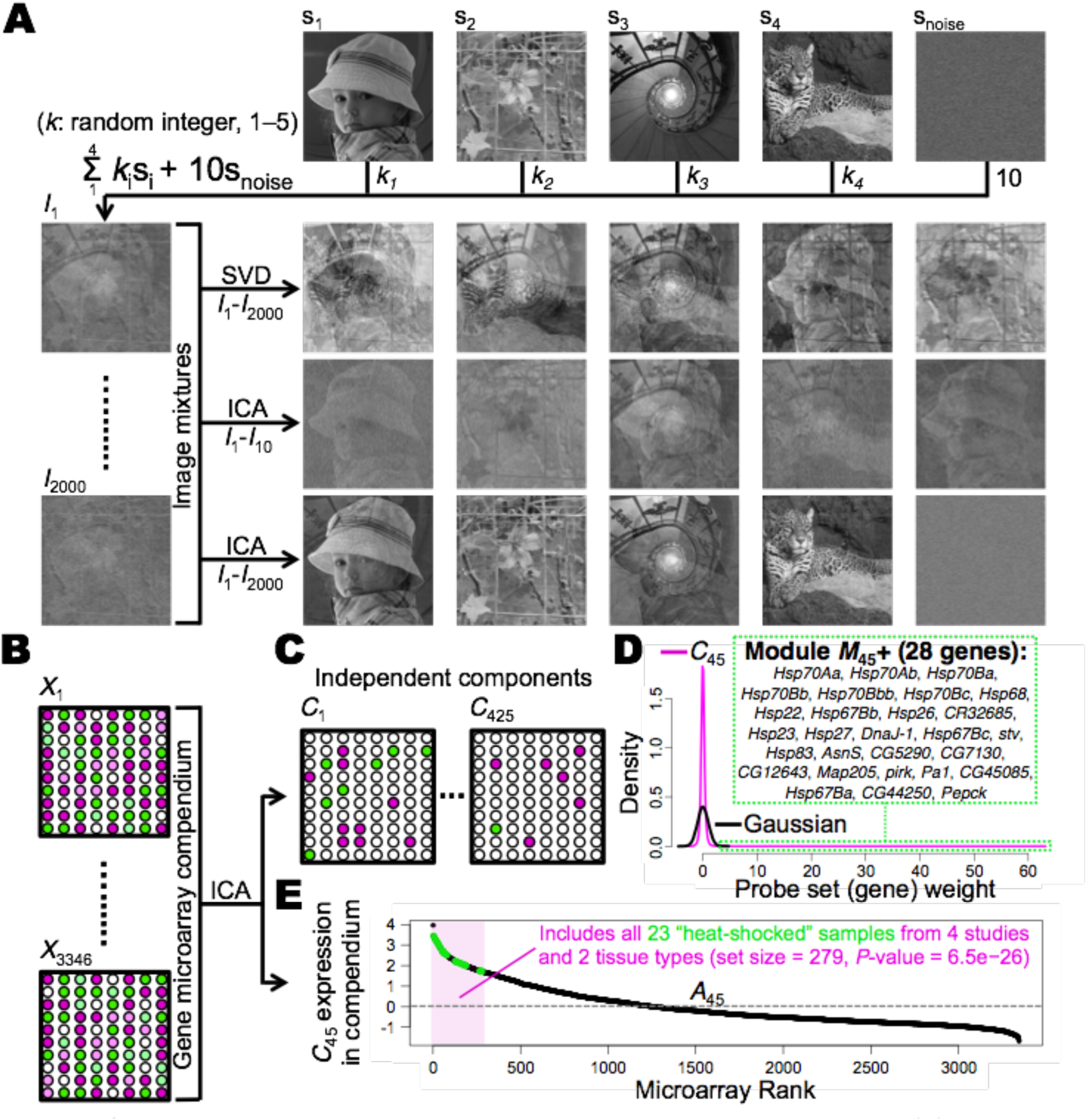
Gene expression module discovery via matrix decomposition methods. (**A**) Independent component analysis (ICA) outperforms PCA achieved by the singular value decomposition (SVD) in estimating source grayscale images (s_1_–s_4_) when given abundant image-mixture data (bottom row). Random coefficients (*k* = 1–5) applied to the sources’ pixel values provide the richness in the image-mixture compendium (*I*_1_–*I*_2000_) for ICA to fit maximally statistically independent pixel intensity patterns. (**B**) Similar to image-mixture pixel values, gene expression levels are a mixture of independent sources of gene expression influences with varying amplitudes and polarity in the microarray compendium (*X*_1_– *X*_3346_). (**C**) We extracted 425 fundamental components of gene expression (*C*_1_–*C*_425_) using ICA on microarrays from diverse *Drosophila* samples and experimental conditions. Unlike the picture components affecting all pixels, biological components mostly impact a relatively small number of genes and so have a super-Gaussian distribution with large tails containing the heavily-weighted, most important genes. (**D**) An example of the hallmark super-Gaussian probe set weight distribution of biological ICA components (*C*_45_) plotted with a reference Gaussian. Most gene probe sets (∼18,928) have near-zero weight, but the group with the largest weight (> 3), i.e. the positive gene module *M*_45_+, is highly enriched for known heat shock genes. (**E**) The second output of ICA, the ***A*** or mixing matrix, contains each array’s component expression levels. Shown is the ***A*** matrix profile for component *C*_45_ (*A*_45_). Arrays with high component expression in the ***A*** matrix profile distribution are often enriched for similar experimental conditions. The hypergeometric *P*-value was from testing X ≥ 23 successes (all “heat-shocked” arrays) in an array sample set = 279, as depicted.

For mining gene expression modules with ICA (reviewed in Kong et al., 2008), we downloaded Drosophila Genome 2.0 (Affymetrix) microarray raw gene expression data available from the NCBI Gene Expression Omnibus (Edgar et al., 2002). We only retained data from *Drosophila melanogaster* mRNA measurements that passed the quality control metrics outlined in the *simpleaffy* package (Wilson and Miller, 2005) (see **Materials and methods**). We used the FastICA algorithm on 3,346 high quality arrays from 221 studies constituting our compendium after preprocessing (**Figure 1B**) and extracted an optimized set of 425 fundamental components of fly gene expression influence (**Figure 1C; Supplementary file 1**). Optimization of the module set is described in detail in the Materials and methods section. Unlike SVD singular values, FastICA independent components are outputted in no priority order and form a set of components whose biological importance must be ascertained downstream. An example independent component, *C*_45_, exhibits a super-Gaussian distribution of gene microarray probe set weights that confer the degree of membership for each probe set in the expression pattern (**Figure 1D**).

Super-Gaussian distributions are hallmarks of biological ICA components found in other species from yeast to human (Kong et al., 2008; Lee and Batzoglou, 2003; Liebermeister, 2002). For such distributions the majority of gene weights are near zero indicating limited involvement of most genes in any particular transcriptomic signature. Super-Gaussian distributions have large tails of high absolute gene weight (three standard deviations away from the mean; the criteria used in Lee and Batzoglou, 2003, and in Engreitz et al., 2010) that discretize relatively small groups of genes—gene modules—that are the most important members of the signature. For example, *C*45’s positive gene module (Q*M*_45_+) contains only 28 genes corresponding to 25 out of 18,952 gene probe sets of the microarray platform used in our compendium (**Figure 1D**). This small module contains the hallmark heat shock genes and is greatly enriched for DAVID Functional Annotation Gene Ontology (GO) categories (Dennis et al., 2003; Huang et al., 2009) related to the heat shock stress response, including: *Stress response* (FDR < 1.0e−20), *Response to heat* (FDR < 1.0e−18) and *Alpha crystallin/Heat shock protein* (FDR < 1.0e−6). Importantly, poorly described genes present in unbiasedly-generated ICA modules, such as *CG12643* and *CR32865* in *M*_45_+, are therefore candidate genes involved in the enriched processes of the well-described genes in the modules. Thus, *Drosophila* gene expression ICA components are mathematical constructs that delineate biologically-relevant modules of co-regulated genes and can provide immediate hypotheses regarding novel gene function. We have isolated modules enriched for many biological processes including, but not limited to: the cell cycle, mitochondria and energy metabolism, immune response, xenobiotic protection, vesicle trafficking, nervous system function, circadian rhythms and developmental pathways (**Supplementary file 1**).

A second output of ICA data matrix decomposition is the mixing, amplitude or ***A*** matrix (Hyvärinen and Oja, 2000) describing the activity levels of the 425 independent components in each of the assortment of experimental conditions and tissue types that constitute the input microarray compendium (**Supplementary file 2**). *C*_45_’s ***A*** matrix expression profile (*A*_45_) across the microarray compendium reveals that samples explicitly labeled as having been “heat shocked” highly express *C*_45_, and thus, highly express the *M*_45_+ module (**Figure 1E**). All 23 arrays for samples dissected 3–5 h after exposure to 37°C from four different studies on two tissue types are present in the top 279 arrays of *A*_45_ (hypergeometric *P*-value = 6.5e−26). Samples expected to express *C*_45_ are highly ranked in *A*_45_, and so other highly ranked samples with their associated perturbation(s) can be hypothesized to also trigger this heat shock response. In summary, ICA can be used for unbiased gene expression module mining with the added benefit to hypothesis generation through ***A*** matrix insights, i.e. through interpretation of the experimental conditions that most express modules of interest. Each of the 425 components’ discretized gene modules (850 in total from all pairs of component tails) with their associated GO category enrichments, along with basic parameters for independent component and ***A*** matrix distributions, are in **Supplementary file 3**.

### The opposing gene modules of one independent component can correspond to two mutually-exclusive expression signatures of the same tissue

The heat shock related module *M*_45_+ of component *C*_45_ (**Figure 1C**) is relatively rare amongst the module set in that it has no opposing counterpart, meaning no group of negatively-weighted gene probe sets surpass our chosen discretization threshold of −3 (Engreitz et al., 2010; Lee and Batzoglou, 2003) to form a second, negatively-weighted *C*_45_ module. Thus, by our criteria, *C*_45_ is an asymmetric component and one of four such components in our entire set of 425 components (**Supplementary file 1** and **Supplementary file 3**). Most components are symmetric defining two modules: one set of expressed genes and one set of suppressed genes. Component *C*_145_ contains all the well-described brain circadian rhythms "evening transcripts" genes in its negative module (*M*_145_−) and most of the "daytime transcripts" genes in its positive module (*M*_145_+), thus the component is a model for the well-known circadian transcriptional clock (**Supplementary file 1**; Ozkaya and Rosato, 2012).

Another possibility each module is expressed in separate groups of compendium samples in a mutually exclusive manner. We noticed that the symmetric components *C*_115_ and *C*_145_ fall in to this second category. *C*_115_’s positive module (*M*_115_+) is highly enriched for larval developmental delay genes (e.g. *dilp8*) and GO terms (e.g. "wound healing"), while its negative module (*M*_115_−) is enriched for pupal genes and GO terms (e.g. "Metamorphosis") (**Figure 2A** and data not shown, respectively). We believe that ICA may include mutually exclusive gene modules of two sample populations in one component to maximize the non-Gaussianity of the input compendium data projections (Hyvärinen and Oja, 2000). Shared genes of relatively low absolute weight, and thus presumably shared tissue and life cycle stage, for both sample populations appears to be a requirement for such module mergers. If we were able to extract a higher number of components with ICA, we speculate that such modules would segregate into individual more asymmetric components. In conclusion, antagonistic or mutually exclusive transcriptional responses may be found on opposite ends of an individual independent component.

**Figure 2.**
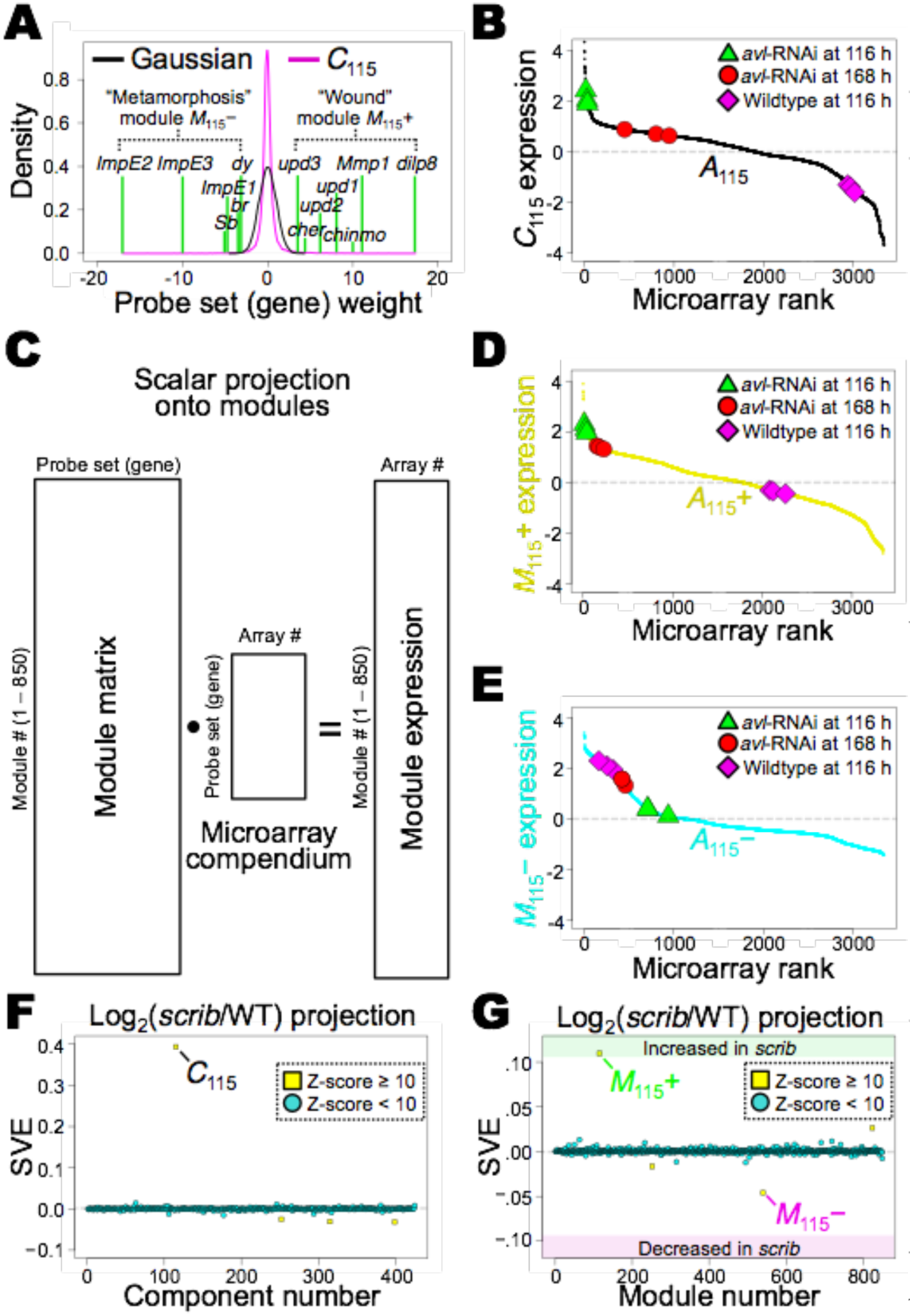
Opposing modules of an ICA component can correspond to two mutually exclusive expression signatures of the same tissue. (**A**) Probability density function of the relatively symmetric component *C*_115_ enriched for wound-healing/neoplastic tumor genes (e.g. *dilp8*) in the positive module (*M*_115_+) and metamorphosis genes (e.g. *ImpE2*) in the negative module (*M*_115_−). (**B**) Distribution of *avalanche* (*avl*) mutant (RNAi) and control third instar larval wing disc samples from study GSE36862 along the *A*_115_ profile. Tumorigenic *avl* mutant wing disc samples at 116 h after egg laying (AEL) have relatively high positive *C*_115_ expression compared to other samples in the compendium and are thus tentatively associated with *M*_115_+ expression. Pupariating control wing discs have relatively negative *C*_115_ expression and are thus tentatively associated with *M*_115_− expression. *avl*-RNAi at 168 h AEL samples known to still express a wound-healing program spearheaded by *dilp8* a n d simultaneously have begun to pupariate at this developmental stage have relatively unremarkable *C*_115_ expression levels. (**C**) We created a module matrix by duplicating the 425 components and then zeroing the weights above and below 3 and −3 in each half of the 850 duplicates, respectively, resulting in 850 module vectors in the module matrix. We then generated a new mixing ***A*** matrix by projecting the microarray compendium onto the module matrix yielding expression profiles for each module across each input microarray. (**D**) Expression level of *M*_115_+ in the microarray compendium, with the samples from **B** marked along the *A*_115_+ activity profile. (**E**) Expression level of *M*_115_− in the microarray compendium, with the samples from **B** marked along the *A*_115_− activity profile. High absolute component *C*_115_ expression levels of *avl*-RNAi at 116 h AEL and control at 116 h AEL samples seen in **B** are confirmed to be mostly due to high expression of the single modules seen in **D** and **E**, respectively. Conversely, mediocre component *C*_115_ expression of *avl*-RNAi at 168 h AEL samples seen in **B** masks the concurrent (entangled) high expression of both opposing modules in these samples shown in **D** and **E**. (**F**) Projection of *scrib*/WT RNA-seq log_2_ gene expression fold-changes onto the 425 components. *C*_115_ explains most of the gene expression variance. (**G**) Projection of *scrib*/WT RNA-seq log_2_ gene expression fold-changes onto the 850 modules. Modules *M*_115_+ (enriched in *scrib*) and *M*_115_− (depleted in *scrib*) explain most of the gene expression variance.

The compendium arrays that express *C*_115_ are largely coherent and include those of wounded, gamma ray irradiated, and neoplastic tumor mutation samples (**Supplementary file 2**). For example, *avalanche* mutant (*avl*-RNAi) larval wing imaginal disc samples 116 h after egg laying (AEL) (Colombani et al., 2012)—known to express the wound/developmental-delay *dilp8*-dependent transcriptional program—highly express *C*_115_ compared to other samples suggesting they highly express module *M*_115_+ and/or are suppressing module *M*_115_− (**Figure 2B**). Conversely, control disc samples from the same study and developmental time point have very low *C*_115_ expression, possibly due to high pupariation module *M*_115_− expression and/or low *M*_115_+ expression. However, we noted that *avl*-RNAi 168 h AEL samples have relatively mediocre *C*_115_ expression levels (**Figure 2B**). Animals of that genotype are still expressing the *dilp8*-dependent transcriptional program in their discs, yet ∼50% of them are pupariating (Colombani et al., 2012). These observations suggested that our model of *Drosophila* gene expression based on the 425 components may fail to resolve complex samples that express *both* opposing modules of a symmetric component because high absolute scalar projection values of opposite sign may cancel out.

To circumvent some of the above shortcomings related to mixed expression samples, we decided to optimize our ICA expression model by basing it on isolated modules (n = 850) rather than whole components (n = 425). We generated a module matrix by first duplicating the independent components and then zeroing weights above and below 3 and −3 in each half of the duplicated component set, respectively. The advantages of this optimized module matrix-based model are that it: 1) allows disentanglement of concurrent expression of two opposing modules in new test data samples that break the general rules for samples in the compendium; 2) solves the issue of sign ambiguity obscuring which part of an expression pattern is upregulated and/or downregulated in projection data by making all weights greater than or equal to zero; and 3) retains the information on which modules originated from the same component with the “+” and “−” nomenclature for tracking their original relationship. We generated a module mixing ***A*** matrix by scalar projection of the microarray compendium onto the module matrix (**Figure 2C**; **Supplementary file 2**; see **Materials and Methods**). No information is lost with the component to module level ICA model transformation.

To demonstrate the utility of the module level ICA model we plotted the module ***A*** matrix expression profiles for the “wound-healing” module *M*_115_+ (*A*_115_+, **Figure 2D**) and “pupariation” module *M*_115_− (*A*_115_−, **Figure 2E**) then marked the same samples from **Figure 2B** along the profiles. We observed relatively high expression of wound-healing *M*_115_+ in both *avl*-RNAi 116 h AEL and *avl*-RNAi 168 h AEL samples and low expression in control 116 h AEL samples (**Figure 2D**). In addition, we observed relatively high expression of pupariation *M*_115_− in both control 116 h AEL and *avl*-RNAi 168 h AEL samples and low expression in *avl*-RNAi 116 h AEL samples (**Figure 2E**). Thus, *avl*-RNAi 168 h AEL samples had more biologically relevant distributions along both module ***A*** matrix profiles (*A*_115_+ and *A*_115_−) compared to their distribution along the original component ***A*** matrix profile (*A*_115_, **Figure 2B**). We further illustrated the utility of the module model by projecting *scrib*/widltype gene expression fold-changes (**Figure 2D,E**; Bunker et al., 2015). We projected these RNA-seq data onto the original 425 components model (**Figure 2F**) and onto the 850 modules model (**Figure 2G**) to quantify component and module expression, respectively. High *C*_115_ expression explains the majority of the variance (SVE ≈ 0.4, **Figure 2F**) and this was revealed to be mostly due to both high *M*_115_+ expression and low *M*_115_− expression (SVE ≈ 0.12 and −0.05, respectively, **Figure 2G**). These results match the biology of *scrib*^1^ mutants used in the study (Bunker et al., 2015) that are characterized by wing disc tumors (unrestricted “wound” module activity) and lethality at the larval stage, i.e. no animals pupariate, and thus, do not express pupal stage genes. Altogether, these results suggest that the 850 ICA modules-based expression model is more useful than the component-based one, at least in these circumstances. Given the benefits of, and the lack of any foreseeable drawbacks to, the module-based model we utilized it in the remainder of this report.

### Purified CNS cell transcriptomes are mixtures of ICA gene expression modules

Our interest in the chemical protection physiologies of the nervous system, particularly those of the blood-brain barrier (BBB), lead us to develop and search for modules related to those processes (Bainton et al., 2005; DeSalvo et al., 2014; Hindle and Bainton, 2014; Hindle et al., 2017; Mayer et al., 2009; Obermeier et al., 2013; Rusan et al., 2014). We searched our ICA modules for the highest probe set weight of *Mdr65,* the gene encoding the primary xenobiotic efflux transporter of *Drosophila* BBB glia (Mayer et al., 2009). We found the positive module of component *C*_94_ (*M*_94_+) to have the maximal *Mdr65* probe set weight, along with high weights for a plethora of other well-described glial genes (**Figure 3A**). This module was highly enriched for GO categories related to BBB physiologies including those related to membrane transport and metabolism (**Figure 3B**). We examined *M*_94_+’s ***A*** matrix expression profile (*A*_94_+) and saw clustering of whole nervous system samples from 28 experiments expressing *M*_94_+ suggesting that it is a nervous system module (**Figure 3C**). Thus, searching for high weights of individual genes of interest in our module resources can quickly reveal putative co-regulated gene patterns or modules of interest with associated experimental conditions and enriched GO terms (**Supplementary files 1,2,3)**.

**Figure 3.**
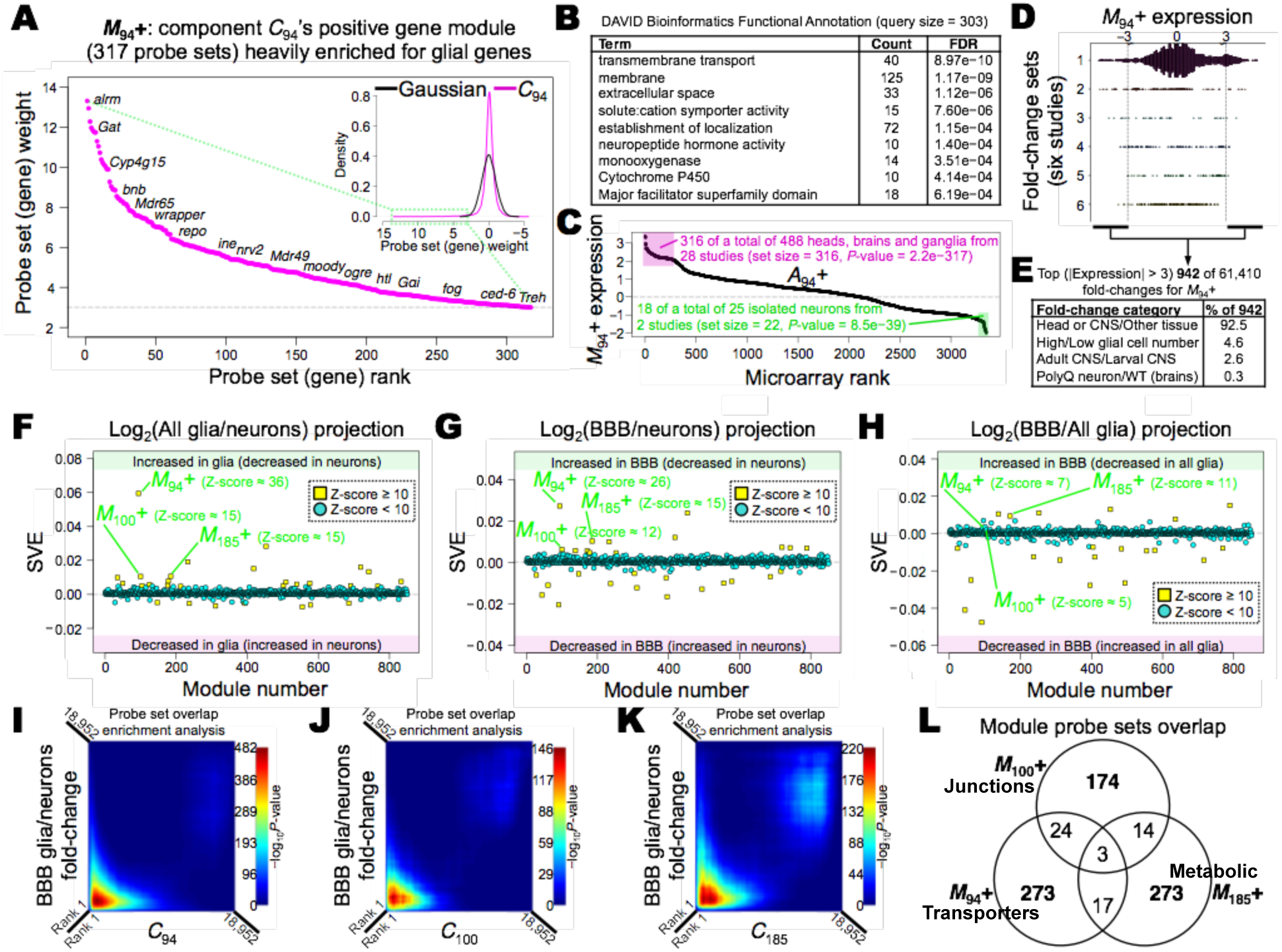
Purified brain cell-type specific transcriptomes are mixtures of concurrently expressed ICA gene modules. (**A**) Positive module of component *C*_94_ (*M*_94_+, 317 probe sets in size) with several of its many known glial genes labeled. (**B**) DAVID Bioinformatics Functional Annotation GO terms highly enriched in *M*_94_+ relate to canonical BBB physiologies. (**C**) The *A*_94_+ profile shows that *M*_94_+ is expressed highly in whole nervous system tissue samples. Isolated neuron samples have the lowest *M*_94_+ expression. (**D**) The top 942 of 61,410 compendium-based gene expression fold-changes in terms of *M*_94_+ expression come from six studies (labeled 1: GSE7763, 2: GSE24917, 3: GSE46317, 4: GSE48852, 5: GSE30627 and 6: GSE27178). Shown are the dot plots of *M*_94_+ expression for all fold-changes from the six studies. (**E**) The top (|*M*_94_+ expression| > 3) 942 compendium-based gene expression fold-changes fall into four categories primarily associated with glial cell mRNA enrichment. Fold-changes with high negative *M*_94_+ expression were inverted to consistently associate a numerator/denominator perturbation pair with those of high increase in *M*_94_+ expression. (**F**–**H**) SVE of log_2_ gene expression fold-change projections onto modules for (**F**) All glia/neurons, (**G**) BBB glia/neurons and (**H**) BBB glia/All glia. *M*_94_+, *M*_100_+ and *M*_185_+ are highly enriched in all glia and significantly more enriched in BBB glia compared to all glia. (**I**–**K**) Rank-rank hypergeometric overlap tests show that the highest ranked gene probe sets in terms of BBB glia/ neurons expression fold-change (i.e. canonical gene expression level enrichment ranking; y-axes) overlap very well with BBB-enriched modules (**I**) *M*_94_+, (**J**) *M*_100_+ and (**K**) *M*_185_+ evident by peak overlap scores occurring in the positive tails of the whole components *C*_94_, *C*_100_ and *C*_185_, respectively, on the left sides of the x-axes (probe sets ranked by positive weight). (**L**) Venn diagram depicting the overlap between gene probe sets of BBB-enriched modules *M*_94_+, *M*_100_+ and *M*_185_+. In conclusion, granular gene expression modules related to xenobiotic and endobiotic transport, septate junction, and metabolic physiologies with largely non-overlapping gene sets are major constituents of the adult BBB transcriptome.

Although module expression levels in the ***A*** matrix provide insights into experimental conditions evoking modules, these levels may be artificially inflated or deflated due to factors unrelated to the reported experimental descriptions associated with each microarray sample. We attempted to limit confounding study-specific factors in explaining the *A*_94_ nervous system sample clustering such as experimenter-specific influences, microarray processing/preprocessing biases and genetic background effects by performing within-experiment projections. Specifically, we projected every non-self sample-pair gene expression fold-change per compendium study onto module *M*_94_+ (see **Materials and Methods**). Of the resultant 61,410 compendium-based *M*_94_+ projection expression levels (centered and scaled projection scalars), the highest (|Z-score| > 3) 942 were from 6 of 221 compendium studies (**Figure 3D**). We interpreted the experimental perturbation pairs constituting these 942 gene expression fold-changes (inverting the numerator-denominator pairs with negative *M*_94_+ expression when necessary) as falling into four categories: 1) Head or CNS tissue in contrast with other tissues; 2) High glial cell numbers in contrast with low numbers, attributed to experimental perturbations with known effects on glial cell count (Avet-Rochex et al., 2014); 3) Adult CNS in contrast with larval CNS; and 4) head tissue with neurons expressing toxic poly-Q repeats in contrast with wildtype head tissue (**Figure 3E**). These results implicate enrichment of glial cell mRNA correlating with *M*_94_+ expression increase. Correspondingly, we concluded that the *A*_94_ clustering of samples expressing *M*_94_+ is due to their shared property of being nervous tissue that contains glial cells. In summary, study-specific effect normalized projections augment ***A*** matrix insights by connecting modules to experimental perturbation-contrasts rather than just to individual compendium sample labels. The ***A*** matrix insights remain important for experiments devoid of condition contrasts that could accentuate a particular expression module (e.g. heat shock module *M*_45_+ where none of the heat shock experiments in the compendium include room temperature controls; **Figure 1D,E**).

To further test whether *M*_94_+ is a glial transcriptional pattern, we projected FACS isolated CNS cell transcriptome data (DeSalvo et al., 2014)—using samples that were not part of the ICA input compendium used to make the model (13 of 15 glial samples) —onto modules (**Figure 3F–H**). All glia/neurons (**Figure 3F**) and BBB glia/neurons (**Figure 3G**) microarray gene expression fold-change projections revealed that *M*_94_+ explains the majority of the expression variance. The BBB glia/neurons microarray gene expression fold-change projection showed that *M*_94_+ is more enriched in BBB glia compared to all glia, although other modules are more BBB-specific (**Figure 3H**). In conclusion, ICA training data and two test data sets implicate *M*_94_+ as an important *Drosophila* BBB glia enriched expression signature of co-regulated genes.

Data-driven identification of modules of interest provides a relatively unbiased route from sample gene expression data to pertinent physiologies. We deduced from the all glia, BBB glia and neuron projections multiple significantly enriched modules in each cell type (**Figure 3F–H**). Namely, from the top twenty modules for each projection, the consistently enriched modules were: all/BBB glial-enriched (*M*_35_+, *M*_50_+, *M*_94_+, *M*_100_+, *M*_104_+, *M*_176_+, *M*_178_+, *M*_185_+, *M*_192_+, *M*_233_+, *M*_418_+, *M*_28_−, *M*_84_−, *M*_294_−, and *M*_317_−); and neurons-enriched (*M*_28_+, *M*_91_+, *M*_126_+, *M*_147_+, *M*_237_+, *M*_278_+, *M*_304_+, *M*_311_+, *M*_357_+, *M*_393_+, *M*_396_+, *M*_69_−, *M*_130_−, *M*_153_−, and *M*_261_−). We highlighted three BBB glia enriched modules associated with well-described BBB physiologies (*M*_94_+, *M*_100_+ and *M*_185_+) and quantified the degree of overlap between gene probe set rankings based on “classic” BBB glia enrichment (i.e. by BBB glia/neurons fold-change values) and gene probe set rankings based on component weights (**Figure 3I–K**). We focused on component *C*_94_ because of *M*_94_+, *C*_100_ because *M*_100_+ is enriched for canonical BBB junction barrier physiology GO terms (**Figure 3—figure supplement 1A**) and *C*_185_ because *M*_185_+ is enriched for *Drosophila* brain energy metabolism GO terms linked to known BBB glia specific physiologies (**Figure 3—figure supplement 1B**; Volkenhoff et al., 2015). The sets of highest ranked genes corresponding to the positive tails of components *C*_94_, *C*_100_ and *C*_185_, i.e. modules *M*_94_+, *M*_100_+ and *M*_185_+, overlap strongly with the sets of highest ranked genes by BBB glia/neurons fold-change (**Figure 3I–K**). Plotted are the hypergeometric test *P*-values (each colored pixel) for extent of overlap of varying gene probe set sizes, incremented by a step size of 8 along the pairs of ranked lists (**Figure 3I–K**; Plaisier et al., 2010; see rank-rank hypergeometric overlap in **Materials and Methods**). Interestingly, modules *M*_94_+, *M*_100_+ and *M*_185_+ are largely composed of unique groups of genes (**Figure 3L**), mirroring their unique sets of enriched GO terms (**Figure 3B**; **Figure 3—figure supplement 1A,B**). We hypothesize that other modules enriched in BBB glia are linked to other independently-regulated biologic processes resident in those cells. In summary, independent modules mapped to biologic processes disentangled by unbiased ICA data mining on whole tissue microarrays can provide comparable information to cell-type specific transcriptomes, without the need for laborious and disruptive cell isolation protocols. Specialized modules concurrently expressed in a particular cell type provide a better view of the different physiologies operating in that cell type, and in addition, modules spanning cell types may best be estimated through data mining of well-controlled bulk tissue sample compendia. We observed several modules with both known glial and neuronal genes, suggesting cross-cell type coordination was indeed captured by the ICA model (**Supplementary files 1)**.

## Discussion

Numerous studies have focused on revealing the transcriptional elements of biological processes (Hawrylycz et al., 2015; Kong et al., 2008). Here, we use ICA—an unsupervised machine learning method—to uncover independently regulated gene expression modules pertaining to a wide range of *Drosophila* biology. By leveraging the diversity and complexity of a large compendium of high quality microarray data we accurately disentangle the mixtures of transcriptional programs that govern fly gene expression. We show how multiple modules map to concurrent biological processes present at the tissue and purified cell population levels, highlighting ICA model modules strongly associated with brain physiologies that remain poorly understood. Our work provides a valuable resource for the important *Drosophila* research system in the form of a well-annotated, quantitative model of gene expression that can be readily explored or used to model new test microarray or RNA-seq data.

Our efforts join those of others that have shown ICA to excel in unmixing gene expression patterns from samples that span many experiments (Engreitz et al., 2010; Kong et al., 2008). By maximizing the statistical independence between the components extracted from the input expression data, ICA delineates groups of co-regulated genes with minimized inter-group regulation dependencies. ICA succeeds in finding such gene expression modules because their levels are observed to vary independently in many samplings of the same expression mixture—the quantified gene levels from the same fly species using the same microarray technology. We utilize a diverse set of samples with regards to tissue type, life cycle stage and perturbation conditions because: 1) we make no biased assumptions about the exclusivity of gene expression to particular sample types. ICA usually improves with more data, and some modules may be expressed in diverse sample types; 2) ICA can handle finding modules that are expressed in subsets of the input mixture, effectively setting their expression very low in the remaining samples. Meaning, ICA is robust against samples considered outliers in the context of a particular module; and 3) a sample type-enriched module may be defined with the aid of samples depleted of the module. A stochastic iterative ICA algorithm, such as FastICA, may arrive at the neighborhood of gene dimensional space defining such a module more efficiently if the projection of samples onto it exhibits a highly non-Gaussian distribution (Hyvärinen and Oja, 2000, 1997). Our discovery that the glial module *M*_94_+ (**Figure 3A**) is shaped by hundreds of CNS-enriched samples of different types and is expressed when CNS-enriched tissue is contrasted with non-CNS tissue (**Figure 3C,E**) is one validation of our compendium selection. However, we suspect that modules built from tissue or developmental stage specific compendia may reflect more nuanced expression responses important for answering different questions. Regardless of compendium type, utilization of a compendium with quality-controlled expression data gathered using the same platform that is preprocessed adequately is a key factor in ICA module quality seen here and in other studies. We suspect the attractive qualities of the *Drosophila* system including but not limited to isogenized stocks, well-controlled experimental conditions, and freshly prepared bulk intact samples contributed immensely to the quality of the input compendium and resultant fly ICA modules.

The success of discovering genes involved in a biological process has correlated with the essentiality of the genes for the process, and the specificity, robustness and prominence of the process. For example, embryonic lethal genes are necessary for development (Nüsslein-Volhard and Wieschaus, 1980). By the same token, heat shock gene discovery was aided by the ubiquity of the stress response, and the reproducibility of the response and eliciting stimuli. Conversely, non-essential genes involved in a post-developmental process distributed across cell types are more likely to remain elusive. Moreover, implication of such non-essential genes in a process through transcriptional profiling may be difficult if few mRNA molecules are required for their function. The rapid advancements of single-cell sequencing technologies coupled with methods such as ICA that can detect subtle yet consistent patterns in abundant data promise to elucidate many unknown links between genes and biological processes (Poirion et al., 2016; Trapnell et al., 2014). Although we and others have shown the utility of inferring gene-to-process connections from mainly bulk tissue expression data provided by multiple experimenters, we expect improved discovery will come from meta-analysis of massive single-cell data gathered by individual groups. Still, we suspect there is much value from models made with the sophisticated ICA methodology applied to freshly frozen bulk tissue samples. Future advances would entail gathering data of similar resolution and breadth to those from single-cells, but obtained directly from intact tissue *in vivo*.

Enormous genomic, transcriptomic, proteomic, metabolomic and other biological data sets are continually accumulating that require efficient and correct interpretation (Schatz et al., 2010). The levels of the various features measured in these data sets all follow the main theme of this paper: they are mixtures of independent sources of influence. Of course, independence of sources is an approximate relationship given the overall interconnectedness of a biological system (Hyvärinen and Oja, 2000; Kong et al., 2008). In practice, methods that maximize the statistical independence of estimated source signals have succeeded in capturing the essential structure of data, suggesting those sources are fundamental and reproducible. For complex biological processes such as those of the CNS or systemic aging, it behooves researchers to disentangle these processes into more manageable pieces; pieces emerging from the inherent structure of the data. Mutually independent “omic-modules” defined by their feature to input data relationships may prove essential for synthesizing a holistic description of biology (Liu et al., 2016). We struggled to define the role of some of our fly modules in part due to poor documentation of sample preparation procedures and the exact genotypes utilized; deficiencies that could have been attenuated (data not shown). If unsupervised analyses that link groups of measured variables to input data are to be used to maximal effect, it is critical for the meta-data to be as complete as possible.

## Materials and methods

### Microarray and grayscale image compendia generation

All GeneChip Drosophila Genome 2.0 microarray (Affymetrix, Santa Clara, CA) raw data (CEL files) available from the NCBI Gene Expression Omnibus (GEO) (platform ID: GPL1322) on September 2nd, 2015 were downloaded using R (R Core Team, 2014). Of the 3,735 downloaded arrays 39 were discarded because the species tested was not *Drosophila melanogaster*, the measurements were not of RNA, the CEL files were corrupt or the data were from an RNA spike-in study. Seven more arrays were discarded because of low n-values (n of 3 or less) in the study that rendered the experimental conditions under-represented. The remaining 3,689 arrays were subjected to the microarray quality control standards outlined in the *simpleAffy* R package (v 2.40.0, Wilson and Miller, 2005). After discarding 340 arrays that did not meet quality control requirements (**Figure 2—figure supplement 1A–C**), three arrays became the sole representatives of their experimental conditions and were also dropped. The final 3,346 arrays of comparable expression distribution were then processed with one of three different variance stabilization methods RMA (Irizarry et al., 2003), GCRMA (Wu et al., 2004) or PLIER (Therneau and Ballman, 2008). Valid combinations of intra-/inter-experiment (by GEO series number; GSE#) probe background intensity correction and summarization, intra-/inter-experiment quantile normalization and technical bias correction (*bias* R package v 0.06, Eklund and Szallasi, 2008) parameters were used with each of the three stabilization methods, resulting in 32 different preprocessed microarray compendia for use in downstream matrix decomposition algorithms.

For generating the 2,000 8-bit grayscale image-mixtures four 512 × 512 pixel JPEG images, with pixel values ranging from 0 – 1, were added pixel-wise after multiplying a random integer (*k*) from 1 to 5 with each individual picture’s pixel values. 10x Gaussian noise generated *de novo* for each mixture using the R function *rnorm* and the pixel value mean and standard deviation of the four images was also added to each picture-mixture. The pixel values of the 2,000 randomly generated image-mixtures were then scaled to the 0–1 in range so they could be visualized. The image-mixture data was stored in a pixel × image-mixture # matrix by linearizing pixel intensity values to 262,144 (512 × 512) rows. Each of the 18,952 probe set × 3,346 microarray or 262,144 pixel × 2,000 image-mixture compendia were used downstream as input data matrices into singular value decomposition (SVD) or independent component analysis (ICA) algorithms.

### Matrix decomposition

SVD decomposes a data matrix into two sets of uncorrelated basis vectors without any assumptions or restrictions on the distributions of the produced vectors. Specifically, SVD decomposes a matrix ***X*** into ***UΣV***^T^ where, in the case of a gene × array ***X*** matrix, ***U*** is a gene × L (where L = rank(***X***)) orthogonal matrix, ***Σ*** is a diagonal matrix and ***V*** is a array × L orthogonal matrix. ***U*** matrix columns are the eigengenes and ***V*** matrix columns are the eigenarrays as described previously (Alter et al., 2000; Holter et al., 2000). The uncorrelated eigengenes are ordered in the ***U*** matrix by the amount of variance they explain in the original data matrix, with the first being the primary principal component underlying the data. SVD on microarray or image-mixture compendia was performed with the *prcomp* R function. Unlike the *princomp* R function, *prcomp* applies SVD to the centered (mean = 0) and scaled (variance = 1) input matrix and uses the n *−* 1 divisor.

ICA decomposes a data matrix ***X*** into the product of two matrices: an ***S*** matrix of statistically independent vectors, and an ***A*** matrix of mixture vectors defining the amplitudes (encodement) of the ***S*** vectors in the original data. ICA was performed using the fixed-point algorithm FastICA (Hyvärinen and Oja, 1997) implemented in the *fastICA* R package (v 1.2-0, Marchini and Ripley, 2013) included in the *fcanalysis* package (Engreitz et al., 2010). Centering and scaling of the input matrix is performed before a whitening, or sphering step. Whitening in FastICA removes second-order statistical relationships by projecting the data onto the first N principal components of the original data set, i.e. it decorrelates the data (Hyvärinen and Oja, 2000), where N corresponds to the number of components extracted from the data. N was varied from 25 to 450 with a step-size of 25 during optimization of the final component set. N independent components are successfully extracted if the algorithm converges to a set threshold of acceptable maximized *S* matrix non-Gaussianity (we used the default for *fastICA* from the *fcanalysis* package, *tolerance* = 0.0001). All other *fastICA* parameters were left at default settings. In the case of a gene × array ***X*** matrix, ICA generates a gene × N ***S*** matrix and an N × array ***A*** matrix. In summary, both SVD and ICA produce uncorrelated gene expression basis vectors for a microarray data compendium matrix, but the ICA components are also maximally non-Gaussian thus rendering them statistically independent. Statistical independence is a stricter condition than uncorrelatedness enforced by ICA in the generation of independently-regulated gene pattern vectors.

### Optimization of the ICA module set

Due to the stochasticity of ICA and the affect of component number choice on the algorithm’s convergence solution we chose to optimize our module set prior to downstream analyses (**Figure 1—figure supplement 1A–I**). Data preprocessing impacts matrix decomposition success and quality so we first confined preprocessing to arrays with similar expression distributions determined by the quality control steps in the *simpleAffy* R package (**Figure 1—figure supplement 2A–C**). Next, we used three variance stabilization methods on the retained data: RMA (Irizarry et al., 2003), GCRMA (Wu et al., 2004) and PLIER (Therneau and Ballman, 2008). We combined microarray background correction, probe set summarization, inter- and intra-experiment quantile normalization and bias correction (Eklund and Szallasi, 2008) where possible to yield 32 different compendium preprocessing conditions. For each preprocessing condition, we extracted five ICA sets of component size 200 after obtaining FastICA convergences (5– 103 FastICA runs required per condition, **Figure 1—figure supplement 2D)** and retained the first 200 singular vectors of each of the unvarying SVD sets (Alter et al., 2000; Holter et al., 2000). We tested the biological relevance of the equally-sized component sets from the two algorithms for every condition (**Figure 1—figure supplement 1,B,E,F**). We measured kurtosis, or distribution “tailedness” (Hyvärinen and Oja, 2000), of component gene expression distribution and found that ICA finds significantly more super-Gaussian (kurtosis > 3) components with larger tails than SVD (**Figure 1—figure supplement 1A**) meaning ICA models relatively small sets of genes as being co/anti-regulated compared to SVD sets. In addition, ICA gene modules had significantly more GO category enrichment than SVD modules irrespective of the preprocessing condition **(Figure 1—figure supplement 1B)**. Based on these two metrics, ICA decomposition using either RMA or GCRMA preprocessing yielded the most biologically relevant components.

Most interestingly, the ability to classify a well-known disease transcriptional state was best achieved using ICA components (**Figure 1—figure supplement 1C–F**). By modeling the gene expression fold-changes between test and control samples as a mixture of the expression levels of the fundamental ICA components, we can quantify component shares of gene expression variation in new test-control data. This modeling resembles the compendium ***A*** matrix but only concerns a single gene expression fold-change vector, i.e. one experiment (**Figure 1—figure supplement 1C**). Larval imaginal discs mutant for the cell polarity organizers *scribble* (*scrib*) and *discslarge* (*dlg*) are known to phenocopy on multiple levels, including transcriptionally (Bilder and Perrimon, 2000; Bunker et al., 2015), and in concordance their gene expression log_2_ fold-changes have similar ICA component expression profiles (**Figure 1—figure supplement 1D**). Scalar projections of mutant versus wildtype control (WT) imaginal disc RNA-sequencing (RNA-seq) expression log_2_ fold-changes (from Bunker et al., 2015) onto components—after mapping probe sets to gene names for RNA-seq to microarray compatibility—transformed gene expression variation into normalized component expression levels. These normalized expression levels, or signed variance explained (SVE) represent the signed fraction of total variance explained by each component. SVE for each component is determined by L2 normal scalar projection (form of quantitative pattern-matching), and thus, a high absolute SVE equals high absolute log_2_ fold-change values matching high absolute component gene probe set weight values, with the signs being maintained for completeness. We found that SVE correlation analysis revealed up to a 98% transcriptional similarity between *scrib* and *dlg* based on ICA component expression (**Figure 1—figure supplement 1E**), a higher concordance than both SVD component expression (< 76%) and the reported 46-71% similarity based on gene overlap in differential expression analysis (Bunker et al., 2015). Thus, a higher *scrib* and *dlg* phenocopying based on SVE correlation is observed using ICA components. We utilized the metric of phenocopying to optimize the module set further.

To avoid over-fitting our ICA model based on one data set, we also tested WT biological replicates from a different RNA-seq study that utilized a high number of biological replicates (Amaral et al., 2014) and found them similar in terms of the standardized phenotype of ICA component expression levels (**Figure 1—figure supplement 1F**). ICA and SVD performances were comparable regarding all WT/WT versus WT/WT comparison (> 0.94 SVE Pearson correlation coefficients using three different WT samples), with ICA outperforming SVD in all but three preprocessing conditions. We attributed this similarity to successful modeling by SVD and ICA of the differences in expression between conditions that are maximally-controlled for being identical, which should reflect Gaussian noise signals stemming from the biology and transcriptional-profiling methodology. Overall, ICA outperformed SVD in several metrics given any preprocessing condition, with RMA and GCRMA being the best variance stabilization methods.

Having confirmed ICA with RMA or GCRMA as the best matrix decomposition method for our purposes, we sought to optimize the specific preprocessing condition and number of components. We discovered that RMA-based ICA component weight distributions in the ***A*** matrix had the lowest average kurtosis, with intra-experiment (per experiment) probe set background correction and summarization having the lowest of this subset of RMA preprocessing conditions (**Figure 1—figure supplement 1G**). This means that component expression level profiles were more Gaussian, and thus, that the components were shaped by a larger fraction of the input data compared to other component sets—a presumably positive property for a general model of expression analysis. Because RMA with intra-experiment probe set background correction and summarization preprocessed compendia were comparable across all metrics examined, we chose RMA without quantile normalization or bias correction from this subset as our ideal minimally preprocessed compendium.

Finally, to optimize module set size, we repeatedly ran FastICA on our compendium until five component sets were obtained per component number requested (for convergence success rates, see **Figure 1—figure supplement 2E)**. We extracted sets of 25–450 components in size in 25-step increments. FastICA failed to converge when > 450 components were requested after 31 runs. Next, we assayed SVE correlations for *scrib*/WT versus *dlg*/WT (**Figure 1—figure supplement 1H**) and WT/ WT versus WT/WT (**Figure 1—figure supplement 1I**) in these sets as described above. We settled on one of the sets of size 425 because: 1) It had the highest *scrib*/WT versus *dlg*/WT SVE correlation of all sets (**Figure 1—figure supplement 1H**); and 2) It had the highest WT/WT versus WT/WT SVE correlation of all sets 75–450 in size (**Figure 1—figure supplement 1I**). Overall our optimized 425 ICA component set delineating 850 modules was enriched for biologically-relevant features and drew on compendium data to a maximal extent. Data from this set was shown in **Figure 1** and utilized in the remaining sections of this report.

### Projection of data onto modules

Scalar projections of expression data onto components or modules were accomplished by computing the dot product of the data and each component/module vector. A log_2_ gene expression vector, ***x***, such as a perturbation-pair fold-change or an individual RMA preprocessed compendium microarray vector, was multiplied with each component/module weight vector of an entire set comprising a component/module matrix, ***S***. Specifically, ***x***.***S*^T^** (where ***S*^T^** is the transpose of ***S***) was computed to yield a mixing vector (for a perturbation pair fold-change ***x***), ***a***, or a mixing matrix (for the microarray compendium, ***X***), ***A***, that describes the activation level or weight of each component/ module in the data. Thus, scalar projections entailed computing ***x***.***S*^T^** = ***a***, or, ***X***.***S*^T^** = ***A***.

***A*** matrix profiles describing the relative expression level of a particular component/module across the compendium microarrays were plotted as ***a*** vector weights, which were centered and scaled, versus microarray rank with regards to expression level. Signed variance explained (SVE) was calculated from a mixing vector, ***a***, by first normalizing the square of each weight value of ***a*** by the sum-of-squares of all weight values in ***a***, i.e. by first calculating the variance explained for each weight value of ***a***. The original ***a*** weight values signs (positive or negative) were restored in the variance explained vector resulting in SVE.

For projection of gene expression fold-changes from non-Affymetrix Drosophila Genome 2.0 chip data (e.g. RNA-seq data), the component/module matrix was first made compatible with the feature IDs of those data using *BioMart* in R (Durinck et al., 2005). Specifically, Affymetrix Drosophila Genome 2.0 chip gene probe sets of component/module matrices were converted to gene names, which resulted in a convergence of 18,952 probe set features to 13,796 gene name features. Component/ module matrix weights for multiple probe sets mapping to a single gene were averaged then assigned to that gene name in the newly formed compatible matrix.

For performing compendium-based projections, each non-self pair of microarray samples per study (per GEO GSE#) in the preprocessed compendium was used to compute a log_2_ gene expression fold-change. These 61,410 fold-changes were centered and scaled then projected onto modules. Two microarrays, GSM216518 and GSM216519, from GSE8775 were excluded from fold-change calculations because they were found to be identical to two other microarrays with different GSM IDs in that same study. Scalar values from all 61,410 projections onto a module of interest (e.g. *M*_94_+ in this study) were then analyzed for perturbation pair category enrichment by manual inspection.

### Statistics and enrichment analyses

One sample Student’s *t* tests were performed with the R function *t.test* on the ICA vs. SVD component kurtosis, GO category enrichment and signed variance explained correlation comparisons using the invariant SVD values as population means (*t.test* parameter *mu*). *P*-values are displayed directly on figure panels and/or in the text. Kurtosis was calculated using the R function *kurtosis* in the R package *moments* (v 0.14). The measured kurtosis was the so-called “classic kurtosis” where a Gaussian (normal) distribution has a kurtosis = 3 and a super-Gaussian distribution has a kurtosis > 3. Pearson correlation coefficients between signed variance explained profiles were calculated using the R function *cor*.

GO category enrichment analyses were performed with DAVID Bioinformatics Resources (v 6.8 Beta; Dennis et al., 2003; Huang et al., 2009) at default classification stringency using all functional categories enabled and with the R package *GeneAnswers* (Feng et al., 2012) on gene module sets, i.e. on probe set groups with members having a weight value above or below 3 or *−*3, respectively. We set a false-discovery rate adjusted *P*-value (FDR) of 0.01 as a threshold for considering a GO category as significantly enriched unless otherwise noted. For the data in **Figure 1— figure supplement 1B**, the significant biological process, molecular function and cellular component GO categories were pooled for each module and for each merged modules per component. ICA outperformed SVD in terms of GO category significance in for each individual GO category comparison as well (data not shown).

*P*-values from hypergeometric tests used in ***A*** matrix profile array enrichment analyses were computed as P(X ≥ *k*), where *k* is the number of successes in the sample set, i.e. the probability of obtaining *k* successes or more given a population size *N*, population successes *K*, and sample set size *n*. Rank-rank hypergeometric overlap tests and plots were done with the *RRHO* R package using a step size of 8 (Plaisier et al., 2010).

Z-scores reflecting the significance of component SVE were calculated by performing 20,000 randomized expression vector scalar projections (permutation tests). Specifically, log_2_ expression fold-change vectors (between test and control conditions) were randomized with the R function *sample*, shuffling the gene probe set or gene name to expression fold-change relationships whilst maintaining overall fold-change variance and magnitude. Randomized data were then projected onto the independent components and Z-scores were calculated for each component by using the mean and standard deviation of the randomized projection scalar values and the scalar value of the original true expression vector.

## Dedication

We wish to dedicate this work to the life and career of our beloved colleague and friend, Dr. John R. Nambu. Thank you for your friendship and eternal commitment to science and mentorship.

## Acknowledgments

We are grateful to Dr. Pat O’Farrell and his lab for the fruitful discussions and nurturing lab meeting environment. We thank current and past members of the Bainton lab for support and feedback on the manuscript. Finally, we thank Angelica and Rania Rusan for the photographs used in Figure 1A.

## Additional information

### Author contributions

ZMR, Conception and design, software creation, acquisition of data, analysis and interpretation of data, and drafting or revising of the article; MPC, software creation, analysis and interpretation of data; RJB, Conception and design, analysis and interpretation of data, and drafting or revising of the article.

## Additional files

### Supplementary files

- Supplementary file 1: The 425 Drosophila independent components (***S*** matrix; genes × components) containing the 850 modules. Additional information obtained from Affymetrix annotations (Release 36; 13 April 2016) regarding each gene probe set including gene names, gene symbols and associated GO terms are in the left-most columns. Individual component weight columns are the remainders. We recommend sorting entire sheet by component weight columns of interest then examining their positive and negative modules at the top and bottom (the tails of the sorted weight vectors).
- Supplementary file 2: Mixing matrices (***A*** matrix; Arrays × components/modules). The component-based ***A*** matrix is in sheet #1, and the module-based ***A*** matrix is in sheet #2. Additional information obtained from GEO regarding each array including experiment ID and experimental procedures are included in the left-most columns.
- Supplementary file 3: Module summary data (Component × properties matrix) that includes information such as length of each opposing module, genes in the modules, component weight summary statistics and associated ***A*** matrix profiles, and enriched GO terms for each module.

---

**Figure 1—figure supplement 1.**
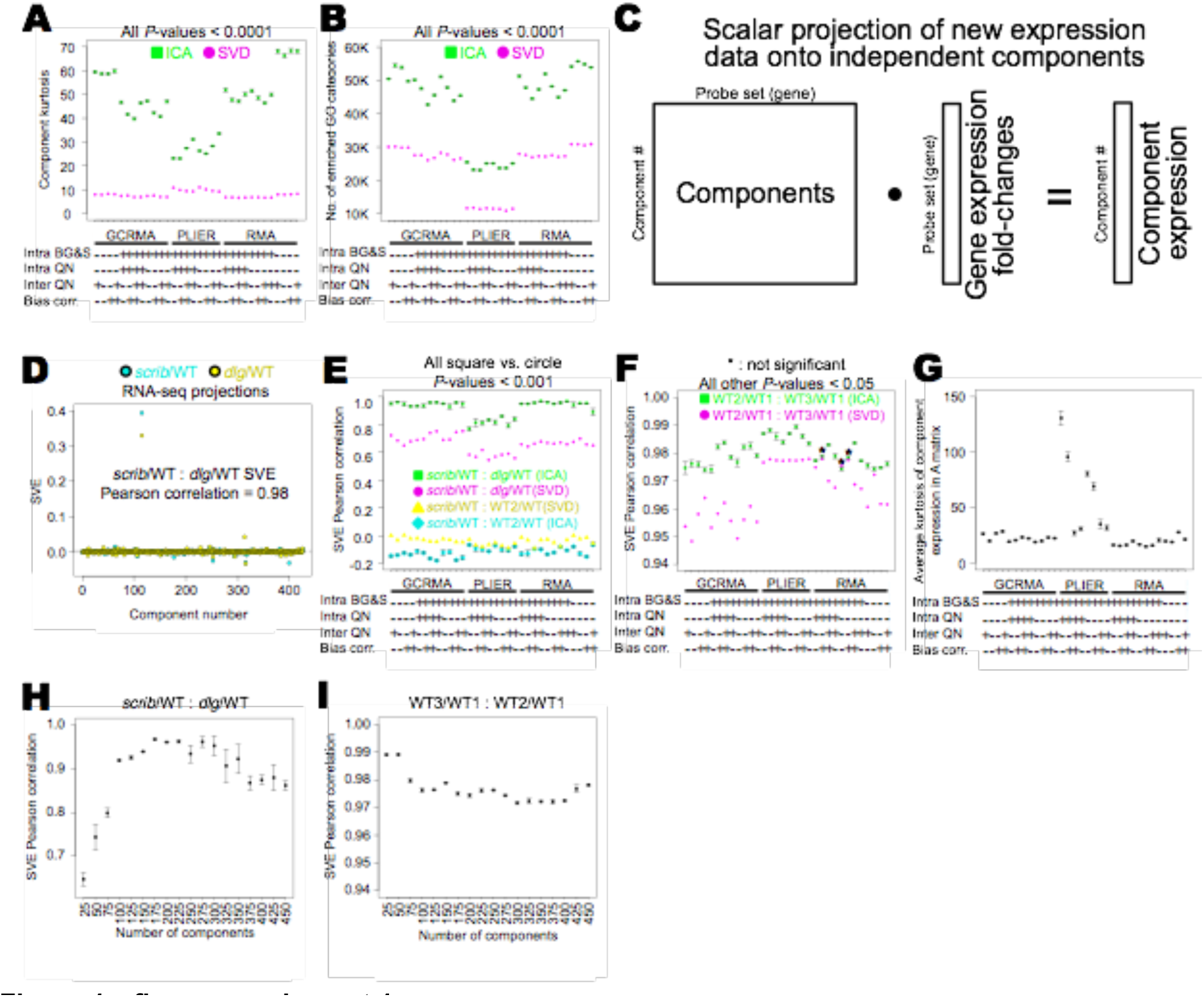
Optimization of ICA components. (**A**–**B**; **E**–**G**) Component sets are of size 200. Preprocessing condition parameters: ‘Intra BG&S’: background correction and probe set summarization per experiment; ‘Intra’ and ‘Inter QN’: per experiment and across experiment quantile normalization, respectively; ‘Bias corr.’: *bias* package technical bias correction; ‘+’ or ‘−‘: method applied or not, respectively. (**A**) Component kurtosis averages of ICA component sets (n = 5 sets per preprocessing condition) are higher than their respective SVD component set averages. (**B**) Highly weighted genes (|weight| > 3; gene modules) of ICA components are more significantly enriched (FDR < 0.01) for Gene Ontology (GO) terms compared to SVD components. (**C**) Scalar projections of log_2_ gene expression fold-changes between test and control samples onto components reveals differentially expressed components in test data (e.g. from RNA-seq). (**D**) Example ICA set projections of gene expression fold-changes for two mutants known to phenocopy (*scrib* and *dlg*) relative to their respective wildtype (WT) controls (i.e. *mutant*/WT). Assuming component activities are a complete model of expression variance (i.e. sum to 100% of variance), we calculated a normalized metric of signed variance explained (SVE) to portray relative component activity levels. A high degree of concordance exists between *scrib*/WT and *dlg*/WT component activities (SVE Pearson correlation coefficient = 0.98). (**E**) ICA components better reflect phenocopying conditions compared to SVD, evident from higher correlations between *scrib*/WT and *dlg*/WT projections. *scrib*/WT compared to WT2/WT projections (two different wildtype samples from the same study) were dissimilar. (**F**) Using another RNA-seq study, two projections using three wildtype samples (WT1, 2 and 3) were found to match using either ICA or SVD components. (**G**) RMA preprocessed ICA components had ***A*** matrix component expression distributions with the lowest kurtosis, indicating that they were shaped by a broader range of input data. (**H** and **I**) We chose RMA without quantile normalization or bias correction as the optimal preprocessing condition then extracted incremental (step size = 25) component set sizes and measured: (**H**) *scrib*/WT and *dlg*/WT projection correlations as in **D** and **E**; (**I**) Wildtype projections correlations as in **F**. All error bars are ± SE.

**Figure 1—figure supplement 2.**
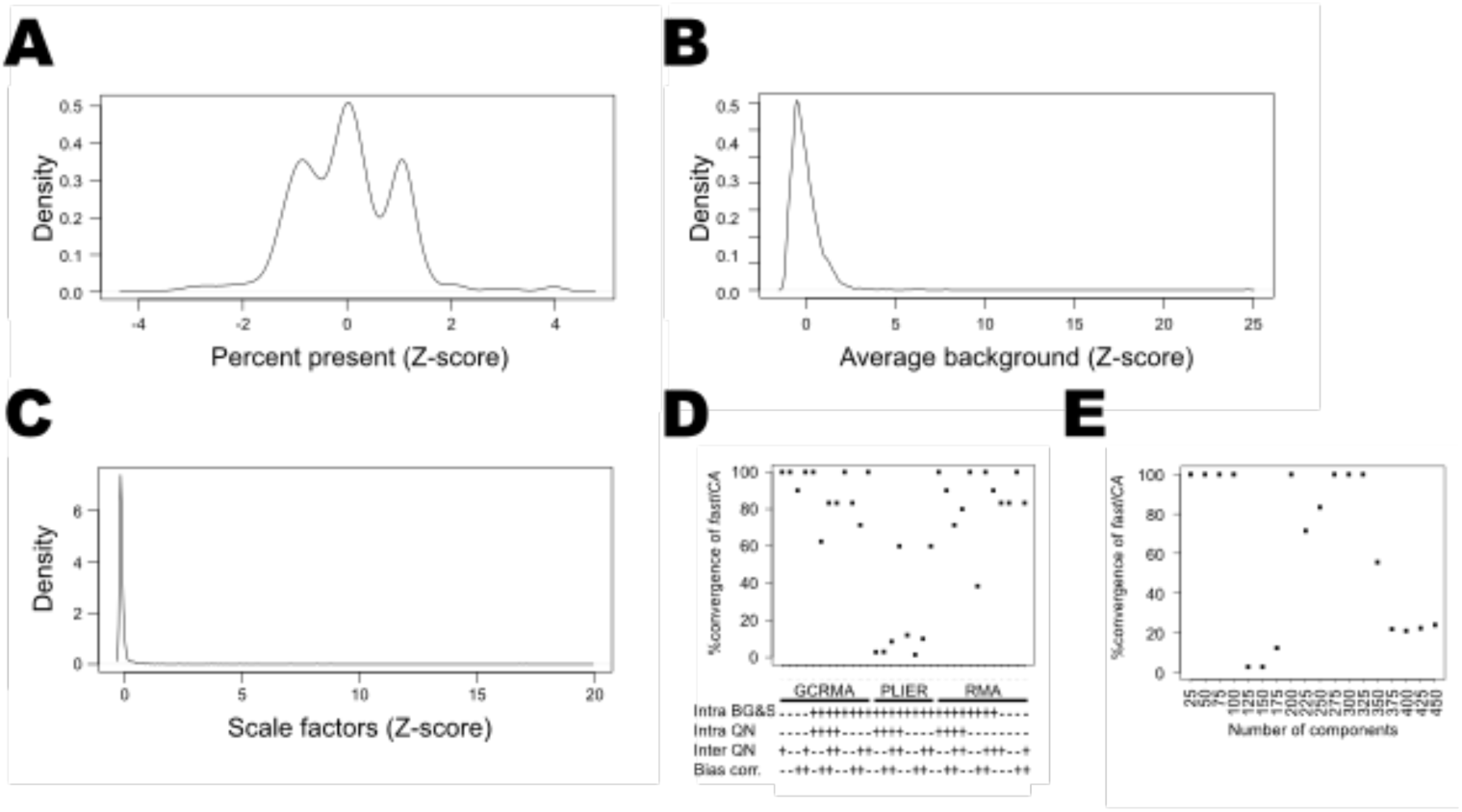
(**A**–**C**) Results from the microarray quality control *simpleAffy* package analysis. Probability density functions for probe set presence (**A**), average background intensity (**B**) and scale factors (**C**) of the downloaded arrays after initial culling of non-mRNA, non-*melanogaster*, spike-in and corrupt arrays. Outlier arrays with absolute Z-scores > 3 for any of the three metrics were discarded resulting in 3,346 retained arrays. (**D**) The %convergence of the FastICA algorithm for each of the 32 preprocessing conditions. The number of FastICA runs needed to obtain five convergences ranged from 5–103, with PLIER preprocessing conditions having the poorest success rate (minimum PLIER condition success rate = 4.9%; n = 103). (**E**) The %convergence of the FastICA algorithm for different component size extractions from the RMA without any quantile normalization or bias correction preprocessed compendium.

**Figure 3—figure supplement 1.**
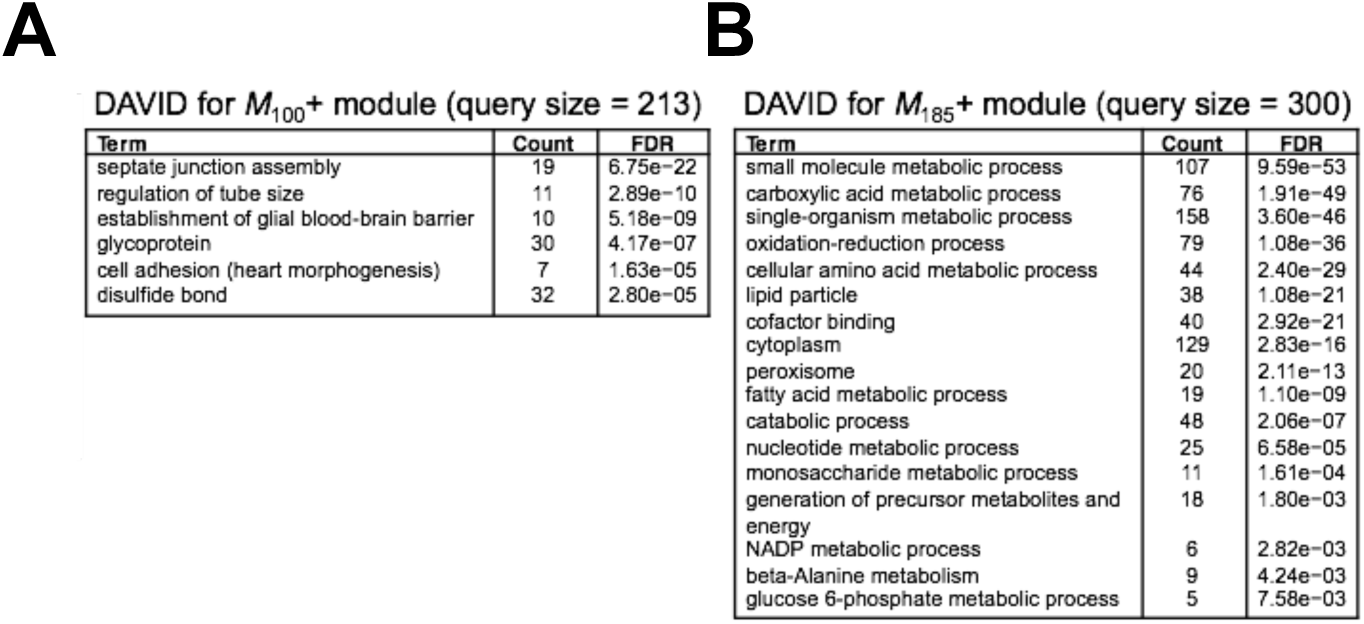
(**A** and **B**) DAVID Bioinformatics Functional Annotation GO terms highly enriched in (**A**) *M*_100_+ and (**B**) *M*_185_+. Separate independent modules with different specializations are expressed in parallel in BBB glia.

## References

Alter, O., Brown, P.O., and Botstein, D. (2000). Singular value decomposition for genome-wide expression data processing and modeling. Proc. Natl. Acad. Sci. U. S. A. 97, 10101–10106.

Amaral, A.J., Brito, F.F., Chobanyan, T., Yoshikawa, S., Yokokura, T., Van Vactor, D., and Gama-Carvalho, M. (2014). Quality assessment and control of tissue specific RNA-seq libraries of Drosophila transgenic RNAi models. Front. Genet. 5, 43.

Avet-Rochex, A., Maierbrugger, K.T., and Bateman, J.M. (2014). Glial enriched gene expression profiling identifies novel factors regulating the proliferation of specific glial subtypes in the Drosophila brain. Gene Expr. Patterns GEP 16, 61–68.

Bainton, R.J., Tsai, L.T.-Y., Schwabe, T., DeSalvo, M., Gaul, U., and Heberlein, U. (2005). moody encodes two GPCRs that regulate cocaine behaviors and blood-brain barrier permeability in Drosophila. Cell 123, 145–156.

Bilder, D., and Perrimon, N. (2000). Localization of apical epithelial determinants by the basolateral PDZ protein Scribble. Nature 403, 676–680.

Brown, J.B., and Celniker, S.E. (2015). Lessons from modENCODE. Annu. Rev. Genomics Hum. Genet. 16, 31–53.

Bunker, B.D., Nellimoottil, T.T., Boileau, R.M., Classen, A.K., and Bilder, D. (2015). The transcriptional response to tumorigenic polarity loss in Drosophila. eLife 4.

Carmona-Saez, P., Pascual-Marqui, R.D., Tirado, F., Carazo, J.M., and Pascual-Montano, A. (2006). Biclustering of gene expression data by Non-smooth Non-negative Matrix Factorization. BMC Bioinformatics 7, 78.

Colombani, J., Andersen, D.S., and Léopold, P. (2012). Secreted peptide Dilp8 coordinates Drosophila tissue growth with developmental timing. Science 336, 582–585.

Comon, P. (1994). Independent component analysis, A new concept? Signal Process. 36, 287–314.

Dennis, G., Sherman, B.T., Hosack, D.A., Yang, J., Gao, W., Lane, H.C., and Lempicki, R.A. (2003). DAVID: Database for Annotation, Visualization, and Integrated Discovery. Genome Biol. 4, P3.

DeSalvo, M.K., Hindle, S.J., Rusan, Z.M., Orng, S., Eddison, M., Halliwill, K., and Bainton, R.J. (2014). The Drosophila surface glia transcriptome: evolutionary conserved blood-brain barrier processes. Front. Neurosci. 8, 346.

Durinck, S., Moreau, Y., Kasprzyk, A., Davis, S., De Moor, B., Brazma, A., and Huber, W. (2005). BioMart and Bioconductor: a powerful link between biological databases and microarray data analysis. Bioinforma. Oxf. Engl. 21, 3439–3440.

Edgar, R., Domrachev, M., and Lash, A.E. (2002). Gene Expression Omnibus: NCBI gene expression and hybridization array data repository. Nucleic Acids Res. 30, 207–210.

Eisen, M.B., Spellman, P.T., Brown, P.O., and Botstein, D. (1998). Cluster analysis and display of genome-wide expression patterns. Proc. Natl. Acad. Sci. U. S. A. 95, 14863–14868.

Eklund, A.C., and Szallasi, Z. (2008). Correction of technical bias in clinical microarray data improves concordance with known biological information. Genome Biol. 9, R26.

Engreitz, J.M., Daigle, B.J., Marshall, J.J., and Altman, R.B. (2010). Independent component analysis: mining microarray data for fundamental human gene expression modules. J. Biomed. Inform. 43, 932–944.

Feng, G., Shaw, P., Rosen, S.T., Lin, S.M., and Kibbe, W.A. (2012). Using the bioconductor GeneAnswers package to interpret gene lists. Methods Mol. Biol. Clifton NJ 802, 101–112.

Hawrylycz, M., Miller, J.A., Menon, V., Feng, D., Dolbeare, T., Guillozet-Bongaarts, A.L., Jegga, A.G., Aronow, B.J., Lee, C.-K., Bernard, A., et al. (2015). Canonical genetic signatures of the adult human brain. Nat. Neurosci. 18, 1832–1844.

Hindle, S.J., and Bainton, R.J. (2014). Barrier mechanisms in the Drosophila blood-brain barrier. Front. Neurosci. 8, 414.

Hindle, S.J., Munji, R.N., Dolghih, E., Gaskins, G., Orng, S., Ishimoto, H., Soung, A., DeSalvo, M., Kitamoto, T., Keiser, M.J., et al. (2017). Evolutionarily conserved roles for blood-brain barrier xenobiotic transporters in endogenous steroid partitioning and behavior. Cell Rep. 21, 1304–1316.

Holter, N.S., Mitra, M., Maritan, A., Cieplak, M., Banavar, J.R., and Fedoroff, N.V. (2000). Fundamental patterns underlying gene expression profiles: simplicity from complexity. Proc. Natl. Acad. Sci. U. S. A. 97, 8409–8414.

Horvath, S., and Dong, J. (2008). Geometric interpretation of gene coexpression network analysis. PLoS Comput. Biol. 4, e1000117.

Huang, D.W., Sherman, B.T., and Lempicki, R.A. (2009). Systematic and integrative analysis of large gene lists using DAVID bioinformatics resources. Nat. Protoc. 4, 44–57.

Hyvärinen, A., and Oja, E. (1997). A Fast Fixed-Point Algorithm for Independent Component Analysis. Neural Comput. 9, 1483–1492.

Hyvärinen, A., and Oja, E. (2000). Independent component analysis: algorithms and applications. Neural Netw. Off. J. Int. Neural Netw. Soc. 13, 411–430.

Irizarry, R.A., Hobbs, B., Collin, F., Beazer-Barclay, Y.D., Antonellis, K.J., Scherf, U., and Speed, T.P. (2003). Exploration, normalization, and summaries of high density oligonucleotide array probe level data. Biostat. Oxf. Engl. 4, 249–264.

Kelley, K.W., and Oldham, M.C. (2015). Transcriptional architecture of the human brain. Nat. Neurosci. 18, 1699–1701.

Kong, W., Vanderburg, C.R., Gunshin, H., Rogers, J.T., and Huang, X. (2008). A review of independent component analysis application to microarray gene expression data. BioTechniques 45, 501–520.

Lee, D.D., and Seung, H.S. (1999). Learning the parts of objects by non-negative matrix factorization. Nature 401, 788–791.

Lee, S.-I., and Batzoglou, S. (2003). Application of independent component analysis to microarrays. Genome Biol. 4, R76.

Liebermeister, W. (2002). Linear modes of gene expression determined by independent component analysis. Bioinforma. Oxf. Engl. 18, 51–60.

Liu, Y., Smirnov, K., Lucio, M., Gougeon, R.D., Alexandre, H., and Schmitt-Kopplin, P. (2016). MetICA: independent component analysis for high-resolution mass-spectrometry based non-targeted metabolomics. BMC Bioinformatics 17, 114.

Marchini, J.L., and Ripley, C.H. and B.D. (2013). fastICA: FastICA Algorithms to perform ICA and Projection Pursuit.

Mayer, F., Mayer, N., Chinn, L., Pinsonneault, R.L., Kroetz, D., and Bainton, R.J. (2009). Evolutionary conservation of vertebrate blood-brain barrier chemoprotective mechanisms in Drosophila. J. Neurosci. Off. J. Soc. Neurosci. 29, 3538–3550.

Nüsslein-Volhard, C., and Wieschaus, E. (1980). Mutations affecting segment number and polarity in Drosophila. Nature 287, 795–801.

Obermeier, B., Daneman, R., and Ransohoff, R.M. (2013). Development, maintenance and disruption of the blood-brain barrier. Nat. Med. 19, 1584–1596.

Ozkaya, O., and Rosato, E. (2012). The circadian clock of the fly: a neurogenetics journey through time. Adv. Genet. 77, 79–123.

Plaisier, S.B., Taschereau, R., Wong, J.A., and Graeber, T.G. (2010). Rank-rank hypergeometric overlap: identification of statistically significant overlap between gene-expression signatures. Nucleic Acids Res. 38, e169.

Poirion, O.B., Zhu, X., Ching, T., and Garmire, L. (2016). Single-Cell Transcriptomics Bioinformatics and Computational Challenges. Front. Genet. 7, 163.

R Core Team (2014). R: A language and environment for statistical computing. R Foundation for Statistical Computing, Vienna, Austria. URL http://www.R-project.org/

Rusan, Z.M., Kingsford, O.A., and Tanouye, M.A. (2014). Modeling glial contributions to seizures and epileptogenesis: cation-chloride cotransporters in Drosophila melanogaster. PloS One 9, e101117.

Schatz, M.C., Langmead, B., and Salzberg, S.L. (2010). Cloud computing and the DNA data race. Nat. Biotechnol. 28, 691–693.

Therneau, T.M., and Ballman, K.V. (2008). What Does PLIER Really Do? Cancer Inform. 6, 423–431.

Trapnell, C., Cacchiarelli, D., Grimsby, J., Pokharel, P., Li, S., Morse, M., Lennon, N.J., Livak, K.J., Mikkelsen, T.S., and Rinn, J.L. (2014). The dynamics and regulators of cell fate decisions are revealed by pseudotemporal ordering of single cells. Nat. Biotechnol. 32, 381–386.

Volkenhoff, A., Weiler, A., Letzel, M., Stehling, M., Klämbt, C., and Schirmeier, S. (2015). Glial Glycolysis Is Essential for Neuronal Survival in Drosophila. Cell Metab. 22, 437–447.

Wilson, C.L., and Miller, C.J. (2005). Simpleaffy: a BioConductor package for Affymetrix Quality Control and data analysis. Bioinforma. Oxf. Engl. 21, 3683–3685.

Wu, Z., Irizarry, R.A., Gentleman, R., Martinez-Murillo, F., and Spencer, F. (2004). A Model-Based Background Adjustment for Oligonucleotide Expression Arrays. J. Am. Stat. Assoc. 99, 909–917.

